# Phenotypic integration and morph-specific strategies in a colour-polymorphic lizard, *Ctenophorus pictus*

**DOI:** 10.64898/2026.05.09.723938

**Authors:** NR LeBas, JL Tomkins, M Olsson

## Abstract

The evolution of alternative male reproductive strategies represents an intriguing evolutionary phenomenon. Divergent strategies are persistently at risk of local extinction or invasion, depending on the suites of traits expressed within and between morphs; hence, understanding the correlational selection that aligns reproductive strategies with behaviour, morphology and physiology is key to understanding the origin and maintenance of genetic polymorphisms. In the polychromatic painted dragon, *Ctenophorus pictus,* yellow, orange and red morphs are well characterised, but the blue morph has been historically absent from studied populations. Here we document the local distribution, morphology and male-contest interactions in a population where blue males are relatively common. We find that blue males express head colouration after a reaching a threshold body size, and that small blue males can reside in close proximity to other males; patterns consistent with a novel size-dependent conditional tactic within the suite of genetic strategies seen in this species. Condition-dependent, positively allometric throat bibs were non-randomly distributed among male morphs, implicating variation in correlational selection and the genetic architecture of the polymorphism. We were unable to definitively assign a morph that was superior in male competition but found that within morphs, male size was the determinant of competitive success, whilst between morphs it was not. Furthermore, contests between morphs were resolved with less aggression than contests within morphs, supporting the idea that badges resolve conflict, and that the invasion of new colour morphs may be facilitated by negative frequency dependent benefits to novel colour variants. These findings highlight the divergent phenotypic, genetic and selective environments that lead to the diversity of colour morphs.

## Introduction

Under evolutionary game theory, alternative strategies can coexist only if they have on average equal fitness through time (Gross, 1996; Shuster & Wade, 1991). In situations where males compete for mates, this is likely through cyclic, frequency dependent fitness pay-offs, such that when a strategy is common it is able to be superseded by one of the alternatives that is rare (Barreto et al., 2017; Maynard-Smith, 1982; Sinervo & Lively, 1996). The children’s game, *rocks-paper-scissors* illustrates the fundamentals of the game, and populations made up of three such unconditional genetic strategies, are expected to exhibit ever changing cycles (Sinervo & Lively, 1996) (but see Corl et al., 2026). In the small number of cases where a genetic polymorphism has arisen in females, these may be highly male-density dependent, as well as frequency dependent (Iserbyt et al., 2013; Sánchez-Guillén et al., 2017).

The general scarcity of examples of evolutionarily stable strategies with three (or more) strategies (Chen et al., 2023; Iserbyt et al., 2013; Kelly, 2024; Pryke & Griffith, 2006; Rankin et al., 2016; Sánchez-Guillén et al., 2017; Shuster & Wade, 1991; Sinervo & Lively, 1996) suggests that the evolution of a third morph is an unusual evolutionary event (Küpper et al., 2016), possibly because demographic or environmental effects limit the persistence of three morphs through space and time (Chelini et al., 2021; McLean & Stuart-Fox, 2014; McLean et al., 2014). Furthermore, once established, very strong correlational selection would be required to maintain the association between phenotypic signal traits and life-history (LeBas et al., 2003) or behavioural traits (Sinervo & Svensson, 2002), ultimately resulting in linkage disequilibrium. Indeed, chromosomal inversions appear to be the origin of at-least some novel morphs, that link and protect (from recombination) a suite of phenotypes together, that is then subject to further selection (Küpper et al., 2016). Nevertheless, evolutionary process promoting novelty in male morphs might arise where new colour morphs are released from competitive interactions because they are not recognized as a threat by rival males. This appears to have occurred in the *Faeder* males in the Ruff (Küpper et al., 2016; Lank et al., 1995), and is proposed for the evolution of colour variation and speciation in cichlid fishes where frequency dependent selection provides a toe-hold for the new colour variant (Seehausen & Schluter, 2004). The evolution of recognition of rival morphs can therefore potentially lead to selection for unrecognised novelty, followed by selection for recognition and so-on in a cyclic manner; a process that could lead to multiple colour signal variants.

Here we investigate the patterns of behavioural dominance within and between morphs in a population containing red, yellow and blue morphs in the colour polymorphic lizard *Ctenophorus pictus*. To date the most intensively studied population of this lizard is at Yathong (NSW), where males are polymorphic – yellow and red morphs are at high frequency with a more scarce orange morph (Olsson, Healey, et al., 2007), however since 1997 blue morphs also established. Life-history trade-offs as well as behavioural variation, maintain stability in genetically polymorphic species. In the Yathong population red males have higher titres of testosterone during their daily activity cycles and yellow males are subordinate to red (Healey et al., 2007; Olsson, Healey, & Astheimer, 2007). Nevertheless, yellow males have relatively larger testes and sire more offspring per-mating (Olsson, Schwartz, et al., 2009). Physiological trade-offs associated with the male morphs has also been documented, with telomere length being longest in yellow males, and shorter in red males, consistent with their higher testosterone, and aggression (Friesen et al., 2017; Rollings et al., 2017). Reactive oxygen species (ROS) production and its relationship with body size follows patterns consistent with the morphs’ life-history strategies (Friesen et al., 2016; Friesen et al., 2020).

In addition to the head colour polymorphism *C. pictus* males are polymorphic for a patch or ‘bib’ of colour on the chest. At Yathong, approximately 40% of red, yellow and orange headed males carry a bib, and bib presence is equally distributed among morphs (Healey & Olsson, 2009). This is reflected more broadly across the species range, where approximately 45% of males in samples greater than 20 had a bib (Baker, unpublished). Within red morphs, bibbed males had higher body condition than non-bibbed males, but the reverse is true in yellow males suggesting that yellow males pay a significant, possibly social cost, for the signal, whereas red males do not. Bibbed males had higher resource holding potential (RHP) and sired more offspring within their territories than non-bibbed males (Olsson, Healey, et al., 2009; Rollings et al., 2017), but had shorter telomeres (Rollings et al., 2017) and higher ROS production (Friesen et al., 2021).

Where organisms engage in contests, the evolution of accurate assessment through rituals can evolve because avoiding escalation and resultant injury when “the battle is already lost” carries a fitness advantage (Parker, 1974). Hence theory expects competitors to know their own resource holding potential (RHP) and accurately assess the RHP of others (Parker, 1974). Where males engage in alternative reproductive tactics, they can be signalled by morphological or chromatic ‘badges’ such as head colour or bib presence or absence. Such badges have been shown to reduce aggression between opponents of recognised RHP (Seehausen & Schluter, 2004), but escalate aggression between matched opponents, e.g. where badges are manipulated (Healey & Olsson, 2008; Pryke, 2007).

In general, data on contest dynamics within and between morphs of species with alternative reproductive tactics remains relatively scarce (Horton et al., 2012; Pryke & Griffith, 2006; Scali et al., 2021). Beyond strategy-denoting badges, some aspects of male phenotype should also signal information about competitive ability that is under selection in encounters within morphs. Hence, where identical strategies meet, outcomes maybe more likely to be based on condition-dependent assessment (Maynard-Smith, 1982; Parker, 1974).

Despite the depth of knowledge on this species, there is a paucity of behavioural and morphological data on the blue morph, due to its rarity in the focal Yathong population., Here, focussed on a South Australian population at Ngarkat, we add to the understanding of aggression and dominance between and within male morphs of *Ctenophorus* pictus in a population where blue males are relatively common.

## Methods

### Collection

Lizards were collected under licence from the Sunset National Park in Victoria, and Billiat and Ngarkat conservation parks in South Australia (Oct/Nov 2002/2003). Specimens were collected by repeatedly driving along a series of dirt tracks and spotting lizards that moved near the track. Any *C. pictus* seen were captured by noosing. The longitude and latitude of the capture point of each lizard was recorded to within 4m using a *Garmin Etrex Venture* handheld GPS. Lizards were placed individually in calico bags and transported to the field station in a cool box to prevent overheating. Captured lizards were maintained outside, in plastic containers (L60 x W35 x H40 cm) with fly-wire lids. Each lizard was housed singly and provided with sand, water and dry eucalyptus bark for shelter. Lizards were fed daily on termites. Cages were moved to full sun in the early mornings/evenings and part shade in the heat of the day. At the completion of the experiments, lizards were returned to their exact site of capture in the field.

### Observations

Trials were conducted with approval of the Animal Ethics Research Committee of the University of St Andrews. Behavioural trials were performed in high sided plastic containers (L60 x W100 x H50 cm). The arenas contained sand and a small amount of eucalyptus bark that one lizard could hide beneath. The arenas were neutral such that each lizard was newly introduced at the time of the trial. Between trials the arenas were washed and the sand and the bark replaced with new material. Observations of behavioural interactions were made from a hide (4WD vehicle). For all the observations, NRL described the behavioural interactions and JLT acted as scribe, documenting times and behaviours. All pairs were size-matched. Male aggression was observed to begin with the raising of the male’s nuchal crest and a stilt-legged display. Often dominance was clearly established without aggression by subordinate males flattening themselves to the ground, while dominant males raised their heads and head-bobbed. Trials were stopped immediately if aggression escalated to biting.

### Morphometrics

The lizards’ snout-vent length (SVL), head depth and head width at its widest point were measured using *Mitotoyo* digital callipers. The weight of the lizard was recorded using portable electronic balance. The head and chest of the lizard were photographed with a Cannon Ixus 330 digital camera. The size of male’s throat bibs were calculated using *image J* image analysis software.

To assess whether body size predicted contest outcome, we used the absolute difference in snout-vent length (SVL) between contestants as a predictor of outcome in logistic regression models. The response variable was binary: whether the larger male won the contest (1) or the smaller male won (0). We fitted separate models for all conclusive contests combined (n=76), within-morph contests only (n=35) and between-morph contests only (n=41), to test whether the importance of size differed between contest types. For blue-red (BR) contests specifically, we used a paired t-test to compare winner and loser SVL directly, to assess whether the tendency for blue males to win against red males could be attributed to a size advantage. All logistic regressions were fitted with a binomial error distribution and logit link function in R. Males were size-matched prior to contests so any size asymmetries reflect imperfect matching rather than deliberate size differences.

### Colourmetrics

The spectral reflectance of the males’ head and chest-bibs were recorded using a spectrophotometer. This analysis is not reported here but supports the assignation of the males to their respective colour morphs.

Statistical analysis was performed in R and SPSS.

## Results

### Snout-vent length and size classes of males

The frequency distribution of male SVL across all populations suggests that there are two size classes of male, and a Shapiro-Wilk test (W = 0.96421, P = 0.007) indicates that SVL is significantly divergent from normal. The nadir in frequency distribution lies at approximately 56 mm (Figure 2). We recorded only a single blue male with a SVL smaller than 56mm (53.7 mm, Figure 3). The ratio of blue males to reds and yellows differed significantly across the two size classes of males, with 6% of blues in the <56mm class, and 36% for reds and yellows combined (χ ^2^_1_ = 6.06, P<0.01). A further consequence of the skewed SVL distribution of blue males is that they were significantly larger than yellows (60.26 +/- 0.75; 56.55 +/- 0.78; Levene’s test, F=4.71, *P* = 0.015) and reds (57.7 +/- 1.17, Levene’s test = F=4.61, *P* = 0.038).

**Figure 1.**
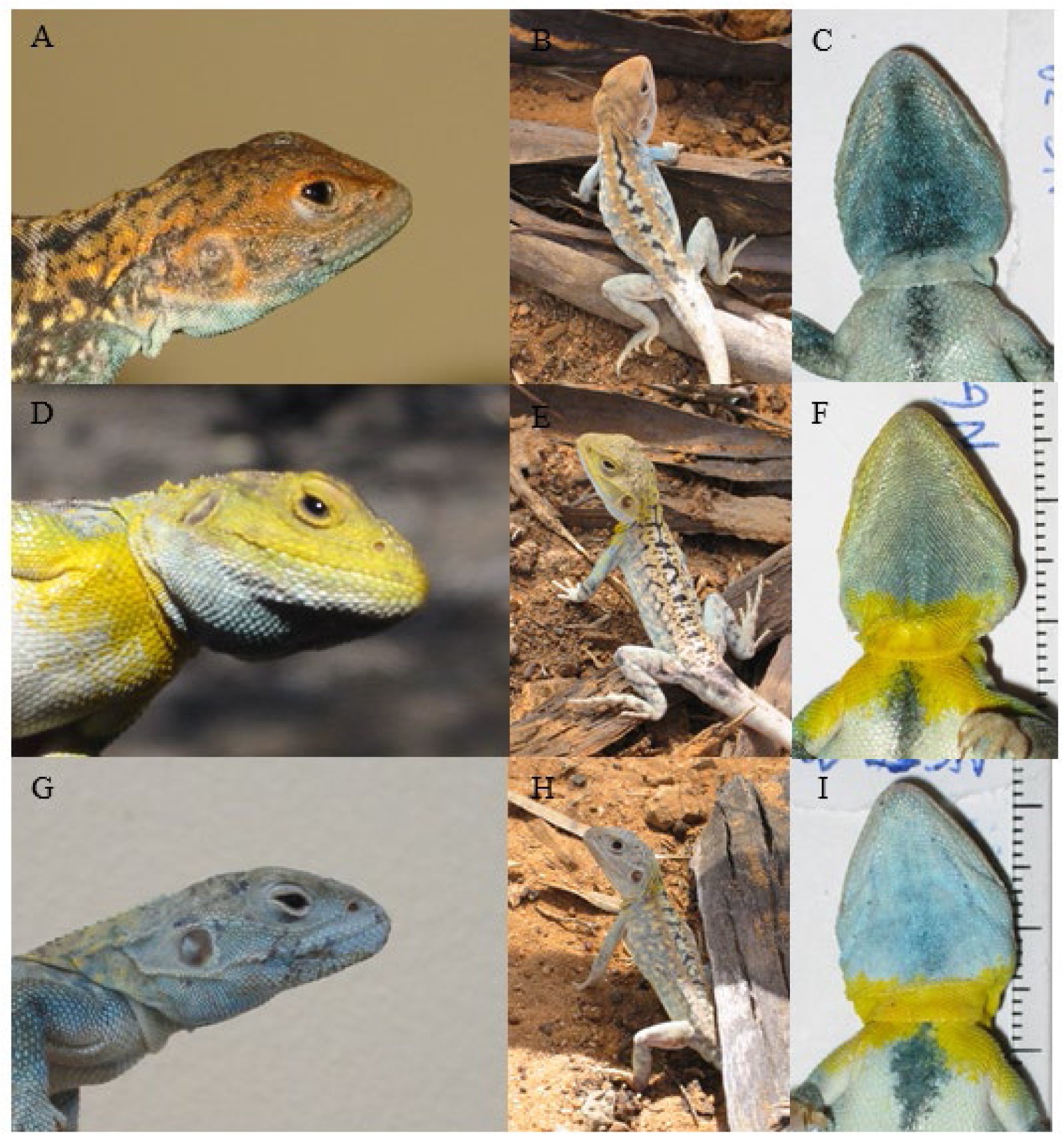
*Ctenophorous pictus* male colour morphs. A, B, C, red headed male with no bib. D, E, F, yellow headed male with bib. G, blue headed male with no bib, H and I Blue headed male with a bib.

**Figure 2.**
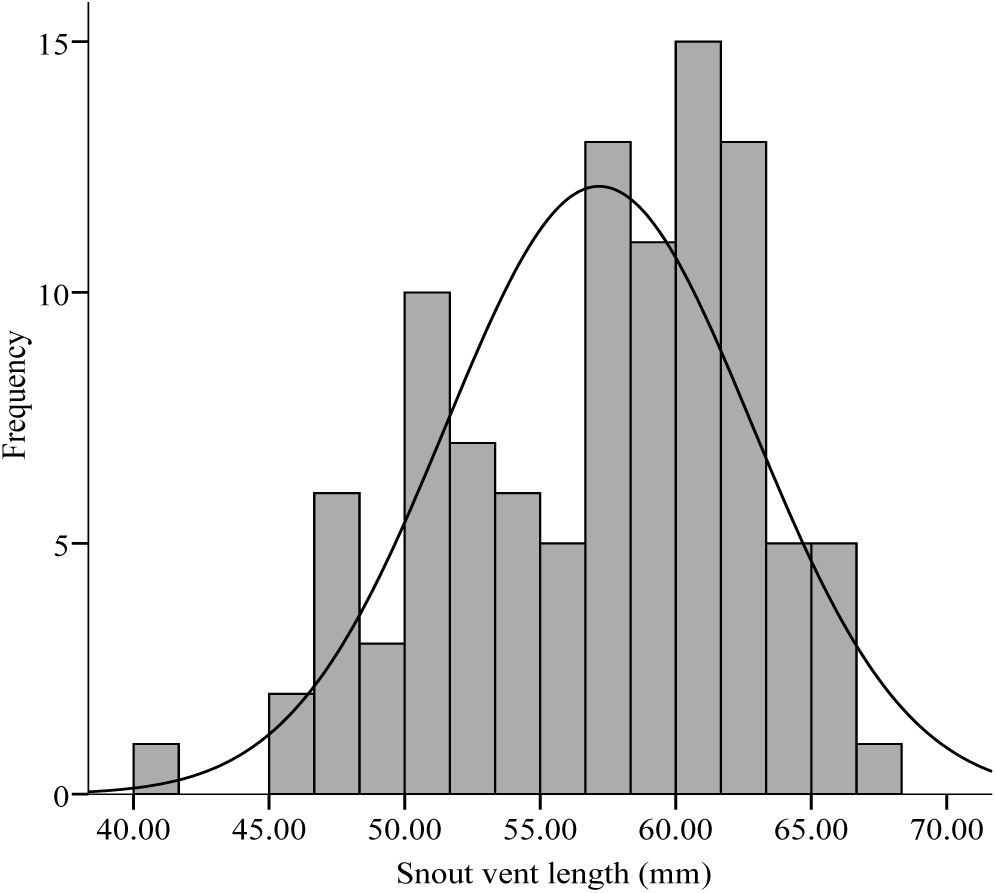
Frequency distribution of snout vent lengths for male *Ctenophorus pictus*, the observed distribution differs significantly from normal (solid line), suggesting that there are two size-classes of males in the sample.

**Figure 3.**
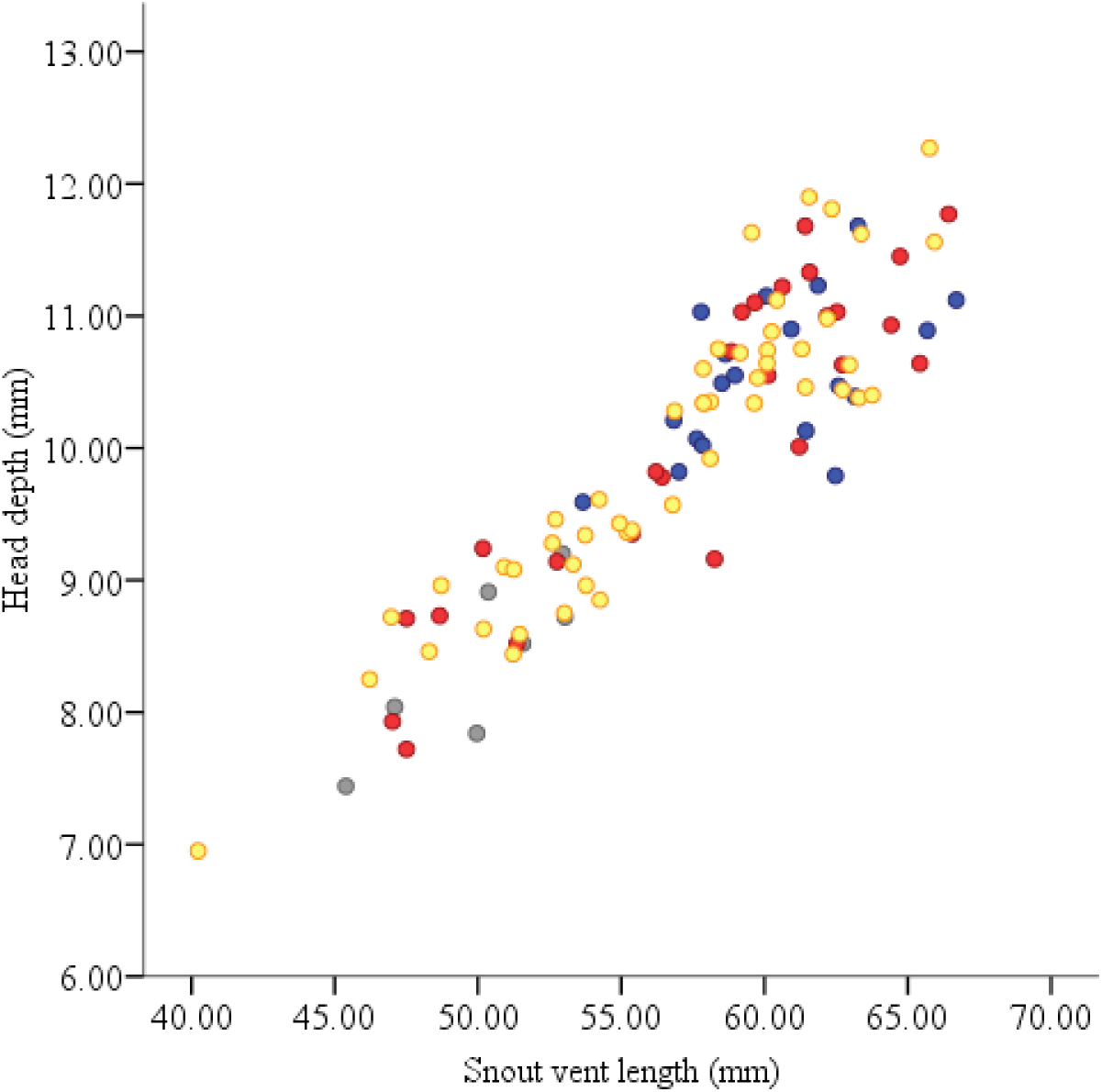
Scatterplot of male head depth on male snout vent length for yellow 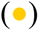, blue 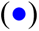 and red 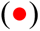 males, and males with no head colour 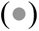. Blue males are almost exclusive to the large size class whereas males with no head colour are present only in the small size class, a pattern that may reflect a developmental strategy.

There are two explanations for the deficit of small blue males, one is that small yellow males (because they are so common in this size class) turn into blues, this would result in a decline in the frequency of yellow males from the small to the large size class. Contrary to this hypothesis however there was an increase in the frequency of yellows from the small (41.5%) to the large size class (58.5%), and these frequencies did not differ from those seen across all males regardless of head colour (35% and 65% respectively; χ ^2^_1_ = 1.33, P>0.1). However, there were seven males for which the head colour was female-like (brown), and all of these males were smaller than 56mm. We hypothesised that these males might be blue-headed, but that the expression of the head colour is delayed relative to the other male morphs and is expressed only when a larger size is reached. Assigning these males as blue accounted for the deficit in small blue males (33% small vs 67% large) and removed the significance of the difference between red and yellow frequencies (36% small vs 64% large) compared to the blues in the small size class (χ ^2^_1_ = 0.04, ns).

### Morph frequencies

Under the paper-scissors-rock game populations are likely to show frequency dependent cycling in morph frequencies since there is unlikely to be a stable ESS where each morph represents a third of the population (Maynard-smith). This means that male morph frequencies should vary in space and time. Hence we have restricted the analysis of male morph ratios to the Ngarkat population for which we have the largest sample: here blue n = 17 (23%), red n = 17 (23%), Yellow n = 41(54%); significantly different from 33.3% (χ ^2^_1_ = 14.3, *P* = 0.0001).

The ratio of bibbed to non bibbed males in yellow and blue head morphs was not significantly different (22 vs 19 and 10 vs 7 respectively; χ ^2^_1_ = 0.004, *P* > 0.05), and was also no different from 50% (χ ^2^_1_ = 0.62, *P* > 0.05). However, there was a significant difference frequency of bibbed to non-bibbed across all three morphs (χ ^2^_4_ = 16.4, *P* = 0.0001). This is due to a deficit of bibbed males in the red morph (0/17) in the Ngarkat population. We only recorded one red male with a bib out of the 26 red males sampled across all populations.

### Distribution

The distances to the nearest male neighbour and nearest female neighbour were calculated from the *Garmin MapSource* software. We estimated that the opportunity to interact with a neighbouring individual would be minimal in lizards captured more than 50m apart, we therefore restricted the following analysis to individuals where the neighbour-of-interest was closer than or equal to 50m. For males and females, a Kruskal Wallis test was used because ‘spread vs level plots’ revealed a relationship between the mean and variance in the data. The nearest other male neighbours were significantly closer for males with no head colouration (χ ^2^_3_= 10.24, *P* = 0.017, Figure 4). Exclusion of the males with no head colour from the analysis reveals that yellow males also have nearer male neighbours than blues and reds (χ ^2^_2_= 6.23, *P* = 0.044, Figure 4). There was no significant variation among the head colour morphs and the distance to the nearest female (≤ 50m away) whether including males with no head colour (χ ^2^_3_= 0.744, *P* = 0.86) or not (χ ^2^_2_= 0.305, *P* = 0.86, Figure 5).

**Figure 4.**
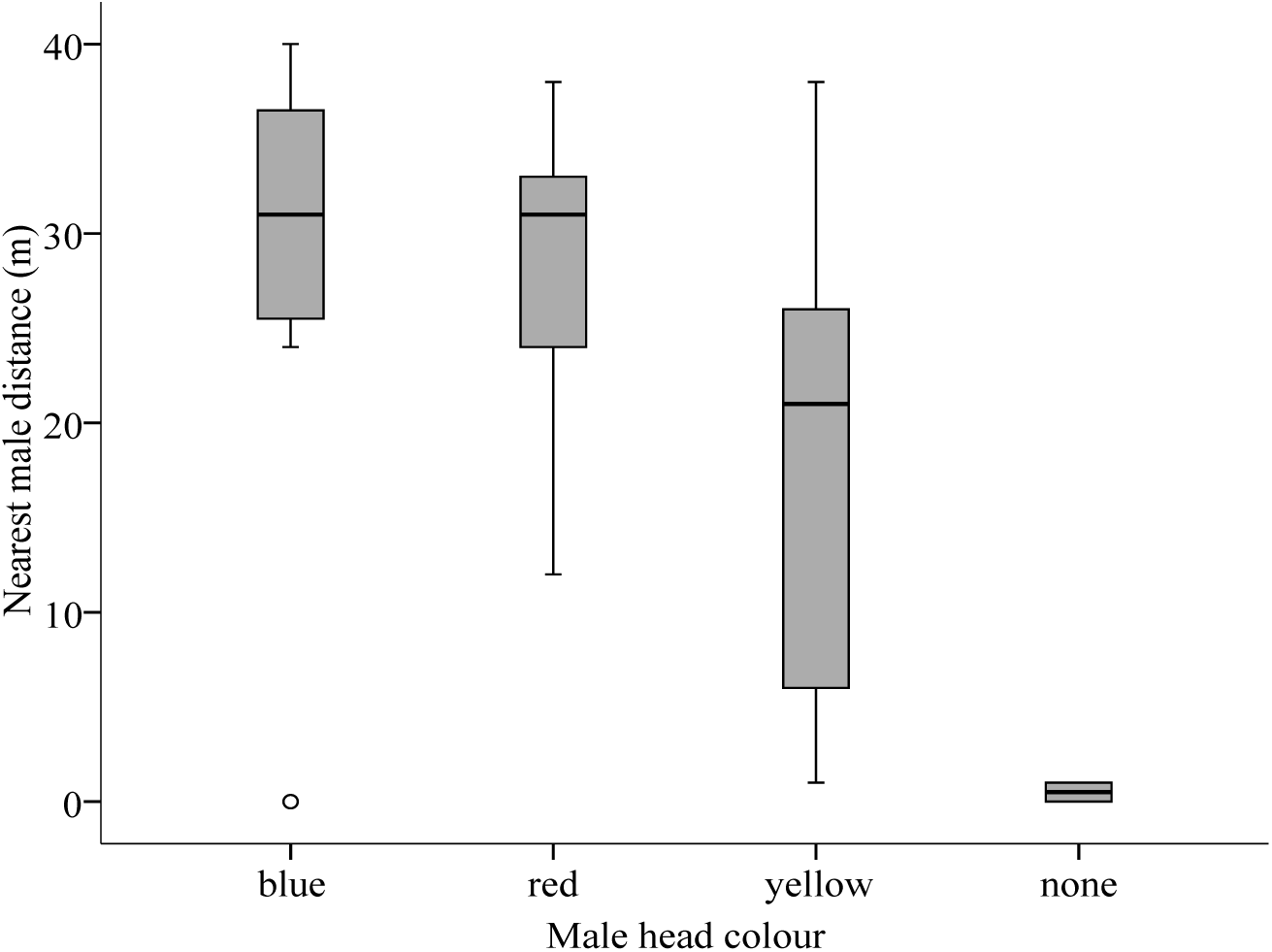
Boxplot (median = line, quartiles = boxes and 95% CI = t-bars) of the distance of the nearest male from each of the male colour morphs and males with no distinguishable head colour. There was significant variation between male head colour morphs and the distance to the nearest male, the significance remains when only blue, red and yellow males are considered.

**Figure 5.**
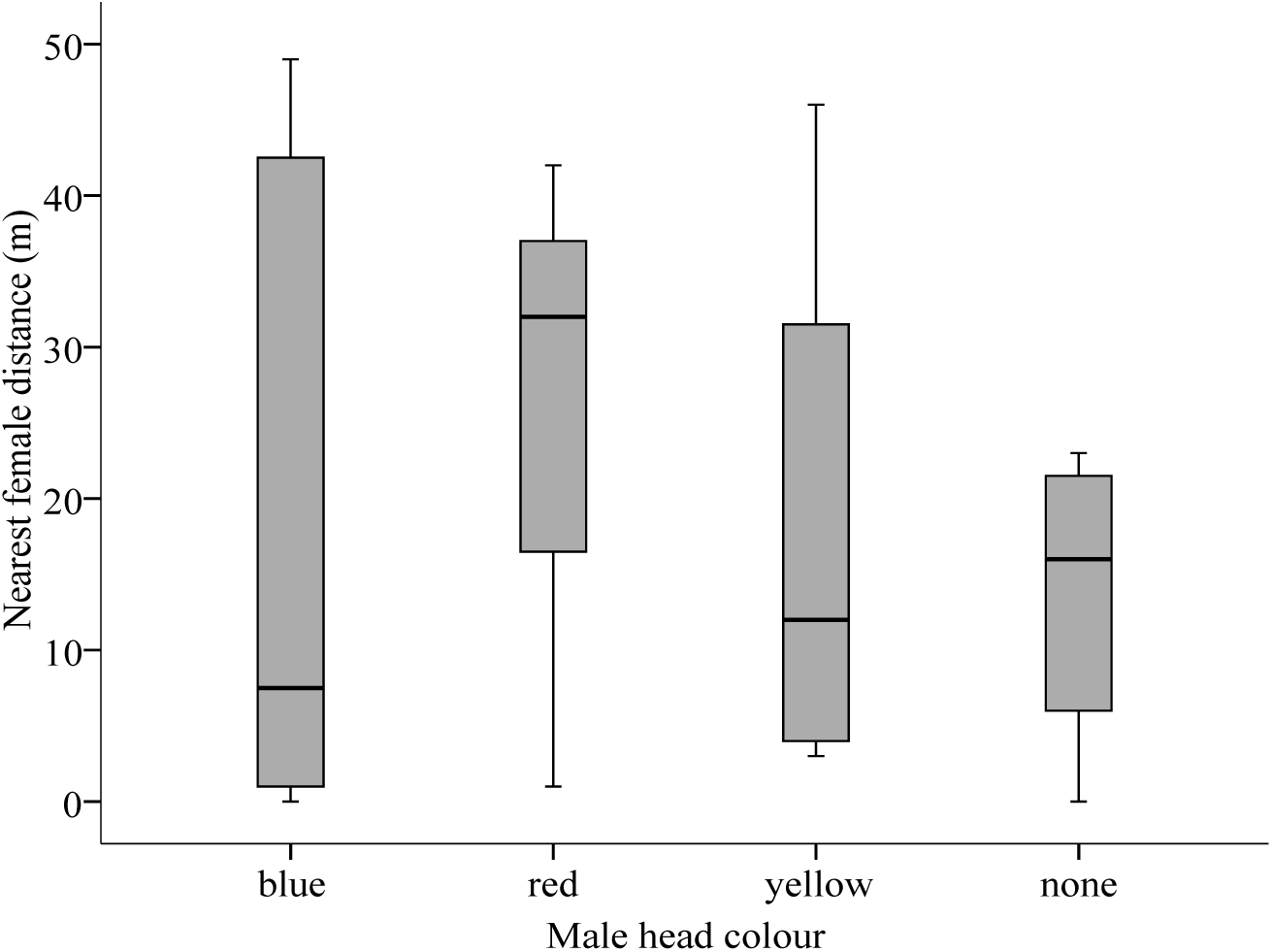
Boxplot (median = line, quartiles = boxes and 95% CI = t-bars) of the distance of the nearest female from each of the male colour morphs and males with no distinguishable head colour. There was no significant variation between male head colour morphs and the distance to the nearest female

Taking each male as a focal individual, the head colour of the nearest male was recorded. This pattern of association was tested against the expected pattern if males assorted randomly with respect to their frequency in the population, so for example we expected 23% of males to have a nearest neighbour that was blue or red and 54% to have a nearest neighbour that was yellow. The two males with no head colour were not included in this analysis because cell counts would be so low. The observed frequencies of head colour morphs varied significantly according to expectation from the population as a whole (χ^2^_4_=13.07, P<0.025, Figure 6). This was primarily due to blue males being more closely associated with reds, and substantially less associated with yellows than expected by chance; the observed pattern of association with other males for focal blue males alone was significantly different from expectation (χ ^2^_1_= 8.78, P<0.005, Figure 6). Both patterns remain significant if the observed frequencies are tested against the frequency of males in the sample (again only distances less than 50m were considered) as opposed to the population as a whole.

**Figure 6.**
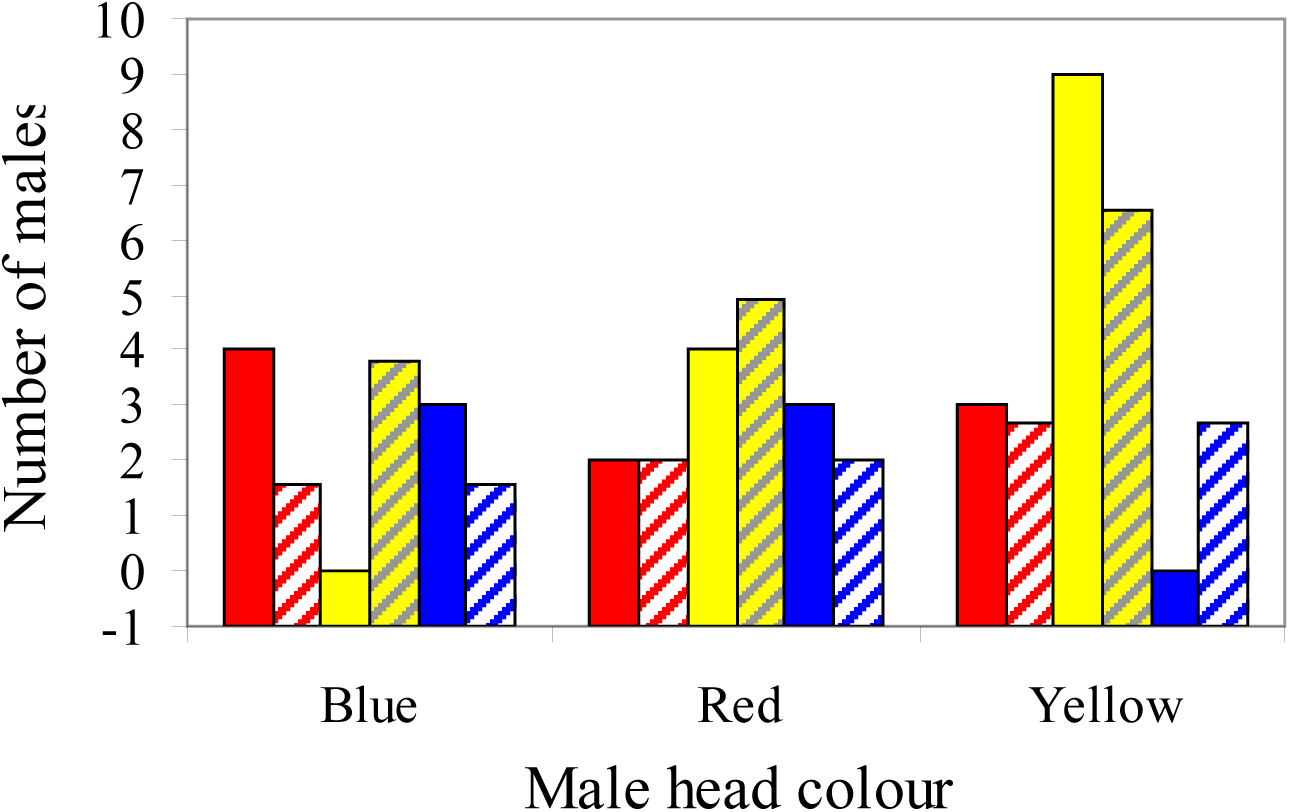
Observed (solid bars) and expected (hatched bars) counts of the nearest male’s head colour to each male when treated as a focal male, focal males are classified by their own head colour. Blue males were more often associated with reds and other blues and less often associated with yellow males, yellow males were more often associated with other yellows.

**Figure 7.**
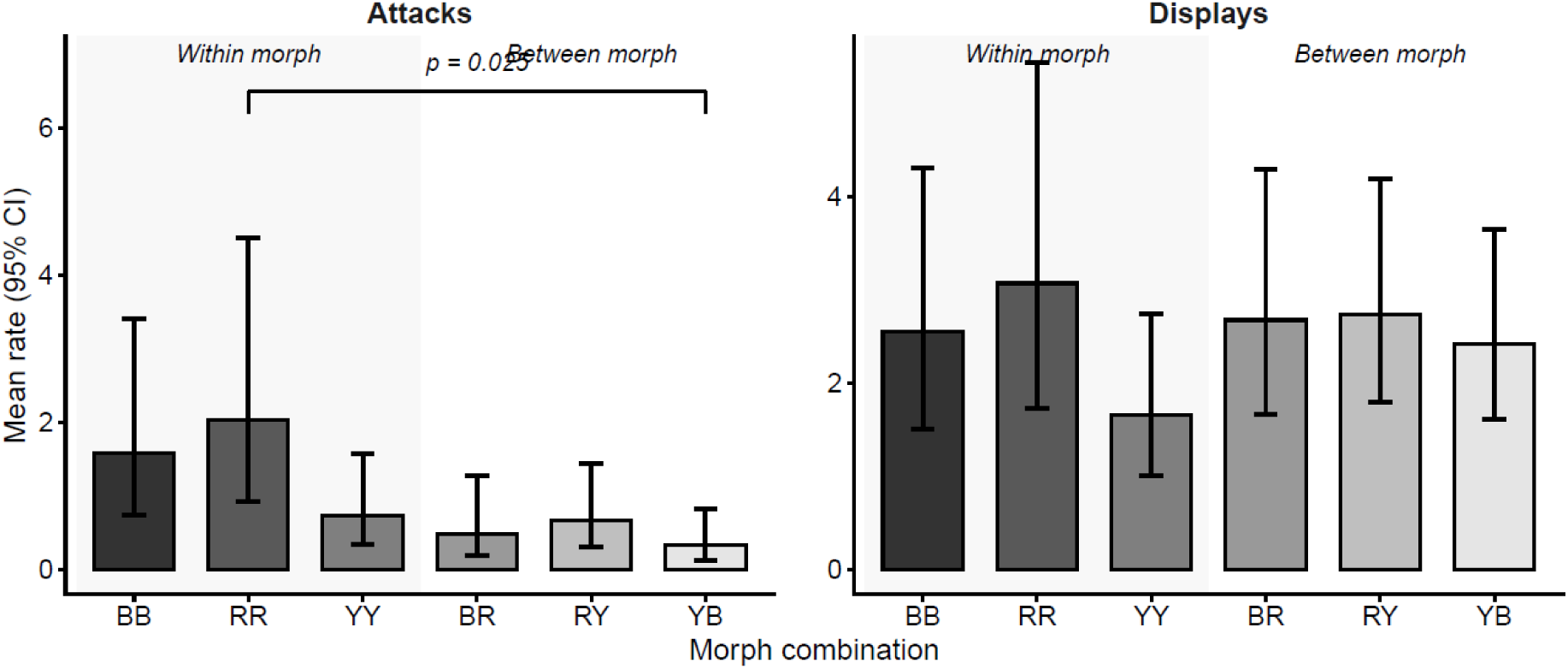
Marginal mean and 95% bars from the GLIM, for the number of fights between all pair-wise combinations of males; pairs of red males tend to illicit the highest levels of aggression.

**Figure 8.**
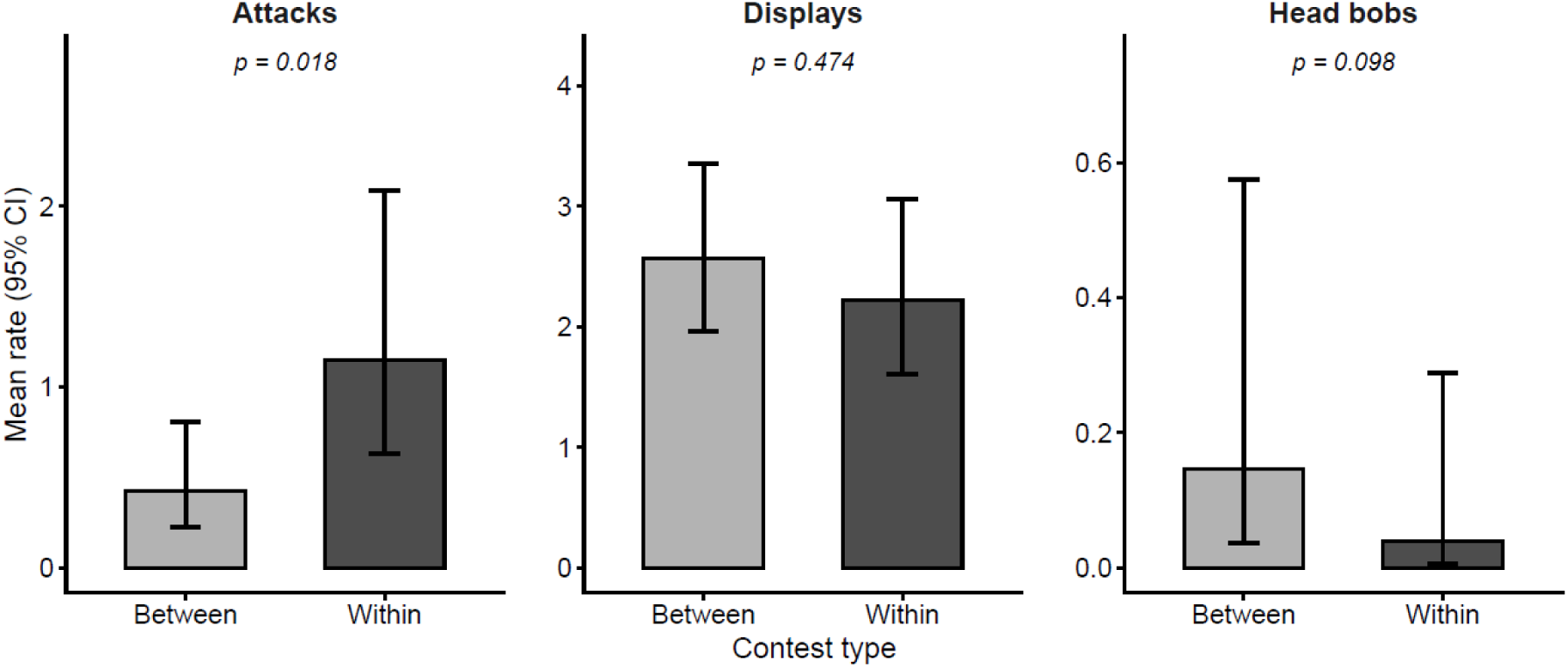
Marginal mean and 95% bars from the GLIM, for the number of fights between morphs or within morphs; same-morph contests illicit the highest levels of aggression.

### Morphology

We used reduced (standardised) major axis (RMA) regression to compare the allometries of head dimensions and areas of the males’ yellow bibs. There was no significant heterogeneity between the red, blue and yellow morphs in the RMA slope or elevation of log head width or log head depth on log SVL, however, the common slope was significantly greater than one for both traits (Table 1). The allometry of weight (condition) did not differ between the morphs (Table 1). All results are robust to the inclusion of small males with ‘body coloured’ like heads into the blue morph category.

**Table 1.** Results of standardised major axis (RMA) regression analyses of head morphology and body mass in *Ctenophorus pictus* males. Section A: comparison among three head colour morphs (blue, red, yellow). Section B: comparison between bibbed and non-bibbed males in blue and yellow morphs only. Slope heterogeneity tests used likelihood ratio statistics. Common slopes and 95% confidence intervals are given where heterogeneity was not significant. Elevation heterogeneity tested using Wald statistic. Mass analyses used log(SVL³) as the measure of body size.

| Analysis | Test | N | Statistic | p |
| --- | --- | --- | --- | --- |
| **Head width ~ SVL** | Slope heterogeneity | 95 | LR = 0.436, df = 2 | 0.804 |
|  | Common slope (95% CI) | 95 | 1.080 (1.017-1.149) | — |
|  | Common slope vs 1 | 95 | F = 7.28 | **0.008** |
|  | Elevation heterogeneity | 95 | Wald = 0.093, df = 2 | 0.955 |
| **Head depth ~ SVL** | Slope heterogeneity | 94 | LR = 0.444, df = 2 | 0.801 |
|  | Common slope (95% CI) | 94 | 1.124 (1.029-1.229) | — |
|  | Common slope vs 1 | 94 | F = 5.76 | **0.018** |
|  | Elevation heterogeneity | 94 | Wald = 0.960, df = 2 | 0.619 |
| **Weight ~ SVL<sup>3</sup>** | Slope heterogeneity | 93 | LR = 0.504, df = 2 | 0.777 |
|  | Common slope (95% CI) | 93 | 1.031 (0.953-1.118) | — |
|  | Common slope vs 1 | 93 | F = 0.92 | 0.339 |
|  | Elevation heterogeneity | 93 | Wald = 0.043, df = 2 | 0.978 |
| **Head width ~ SVL** | Slope heterogeneity | 69 | LR = 0.011, df = 1 | 0.918 |
|  | No bib slope (95% CI) | 69 | 1.084 (0.972-1.208) | — |
|  | Elevation heterogeneity | 69 | Wald = 0.011, df = 1 | **0.915** |
| **Head depth ~ SVL** | Slope heterogeneity | 68 | LR = 4.030, df = 1 | **0.045** |
|  | No bib slope (95% CI) | 68 | 0.942 (0.776-1.143) | — |
|  | Bib slope (95% CI) | 68 | 1.187 (1.053-1.338) | — |
| **Weight ~ SVL<sup>3</sup>** | Slope heterogeneity | 101 | LR = 3.666, df = 1 | 0.056 |
|  | No bib slope (95% CI) | 101 | 0.987 (0.893-1.091) | — |
|  | Bib slope (95% CI) | 101 | 1.118 (1.031-1.213) | — |

Bibbed males occur in the blue and yellow morphs at a frequency of c50%. We tested whether there was an association between male head allometry and expression of the yellow bib in blues and yellows and excluded red males. Since there was no difference in head allometries among morphs, head morph (yellow or blue) was ignored in this analysis. The males with female-like (‘body colour’) heads, two of which had bibs, were also excluded in this analysis. There was no significant variation in the slope or elevation of the head width allometry. However, males with bibs had a steeper head-depth allometry than males without bibs (Table 1). Although, mass increased slightly more with SVL in bibbed males compared to non-bibbed males, the difference was not significant (Table 1). There was no detectable difference in the size of bibbed and non-bibbed males (t_71_= 8.15, *P* = 0.418). The allometric exponent of bib area on log SVL^2^ was significantly greater than 1(Table 2), further bib area was condition-dependent being correlated to residual mass but did not differ between yellow and blue males (Table 2).

**Table 2.** Results of standardised major axis (RMA) regression analyses of bib area and body size and condition males in *Ctenophorus pictus* males.

| Analysis | Contrast | Statistic | P |
| --- | --- | --- | --- |
| **Bib area ~ SVL<sup>2</sup>** | Slope (95% CI) | 3.159 (2.492-4.004) | — |
|  | Slope vs 1 (all bibbed, n = 41) | F = 144.35 | **< 0.001** |
| **Bib area ~ condition** | Pearson r (n = 40) | r = 0.334 | **0.035** |
|  | Yvs B slope heterogeneity | LR = 2.938, df = 1 | 0.087 |

**Table 3.** Results of Poisson mixed models testing the effect of contest type (within vs between morph) on male aggressive behaviour.

| Response | Random effects | Between (95% CI) | Within (95% CI) | LRT $\chi^2$ | df | p |
| --- | --- | --- | --- | --- | --- | --- |
| Attacks | m1.ID + OLRE | 0.43<br>(0.23-0.80) | 1.15<br>(0.63-2.08) | 5.64 | 1 | **0.018** |
| Displays | OLRE only | 2.57<br>(1.96-3.36) | 2.22<br>(1.61-3.06) | 0.51 | 1 | 0.474 |
| Head bobs | OLRE only | 0.15<br>(0.04-0.57) | 0.04<br>(0.01-0.29) | 2.73 | 1 | 0.098 |

### Aggression

#### Contests within and between morphs

Behavioural observations during the trials lead us to assign a winner and a loser in 76 out of 96 cases. Detailed behavioural data were available for 62 of these 77 trials. We qualified our assessment of the winners and losers by analysing the asymmetry in dominant or submissive behaviours across the pairs of males with a Wilcoxon signed rank test. Our assessment of the dominance outcome is supported by the behavioural data; ‘dominant’ males instigated more attacks (Z = 4.62, *P* <0.001) and displays (Z = 6.91, *P* <0.001), while subordinate males exhibited more escape behaviours (Z = 6.21, *P* <0.001). There was a non-significant tendency for winners to exhibit more head-bobs (Z = 1.76, *P* = 0.078).

Male interactions can be classified by the pair-wise combination of male colour morphs, or simply whether colour morphs were interacting within or between morphs. Poisson mixed models testing the effect of contest type (within vs between morph) on male aggressive behaviour (n = 62 conclusive contests). Observation-level random effects (OLRE) were included in all models to account for overdispersion. Male identity (m1.ID) was retained as an additional random effect in the attacks model only, where it explained significant individual variance (LRT χ² = 9.10, df = 1, p = 0.003). Marginal means and 95% confidence intervals are back-transformed from the log scale.

#### Head colour and bib presence

Blue males won more contests against red males, though this did not reach statistical significance (Table 4). No asymmetry was detected between red and yellow, or yellow and blue contests (Table 4). Mixed effects logistic regression was attempted but could not be reliably fitted due to complete separation caused by most individual males appearing in only one or two contests. The Bradley-Terry test for differences in the competitive ability of each morph, no morph was significantly dominant (Table 4).

**Table 4.** Contest outcomes between head colour morphs in *Ctenophorus pictus* (n = 41 between-morph conclusive contests). Win counts and rates are from the perspective of the more frequent winner in each pairing. Binomial exact tests assessed whether win rates differed from 0.5. A Bradley-Terry model estimated relative competitive ability for each morph simultaneously across all pairings (Yellow = 0, reference).

| Contest pairing | Frequent winner | Win count | Win rate | Bradley-Terry | p |
| --- | --- | --- | --- | --- | --- |
| Blue vs Red (BR) | Blue | 9/11 | 0.82 | Blue=0.135 | **0.065** |
| Red vs Yellow (RY) | Yellow | 9/16 | 0.56 | Red=-0.639 | 0.804 |
| Yellow vs Blue (YB) | Yellow | 8/14 | 0.57 | Yellow=0.000 | 0.791 |
| Bradley-Terry overall | — | — | — | LRT $\chi^2=1.95$, df=1 | 0.163 |

When blues and yellows met either within or between morphs, there were sixteen occasions when there was an asymmetry in the presence of the bib, non-bibbed males won 9 times and bibbed males won 7 times (chi2 = 0.25 ns). There was not enough data to determine whether bibbed males were dominant within yellows or blues.

#### Size and fight outcome

We used a logistic regression to test whether males with a larger SVL were more likely to win a contest using the absolute SVL asymmetry between contestants as the predictor. Analyses were conducted for all contests combined, within-morph contests only, and between-morph contests only. For blue-red (BR) contests specifically, a paired t-test compared winner and loser SVL directly. Males were size-matched prior to contests so residual asymmetries reflect imperfect matching rather than deliberate size differences. Across all contests, the larger male was more likely to win (Table 5). However, this effect is driven by within-morph contests, where size was a significant predictor of outcome (Table 5), whereas size did not predict outcome in between-morph contests (Table 5). In BR contests specifically, winners were not significantly larger than losers (Table 5), suggesting that the tendency for blue males to win against red males was not attributable to a size advantage.

**Table 5.** Effect of body size asymmetry on contest outcome in *Ctenophorus pictus*.

| Analysis | N | Larger male won | Test statistic | p |
| --- | --- | --- | --- | --- |
| All contests | 76 | 54/76 (71%) | $z = 2.114$ | **0.035** |
| Within morph only | 35 | 23/35 (66%) | $z = 2.208$ | **0.027** |
| Between morph only | 41 | 31/41 (76%) | $z = 0.519$ | 0.604 |
| BR contests (paired t-test) | 11 | 8/11 (73%) | $t = 1.259, df = 10$ | 0.237 |

## Discussion

Colour polymorphic species represent an opportunity to precisely investigate divergent life-history adaptations, and as such are a powerful tool in evolutionary biology. Here we document the within morph and between morph behavioural interactions and morphology in a population of *Ctenophorus pictus* where the, as yet poorly characterised, blue morph is present. These data highlight how within morph aggression escalates to condition-dependent outcomes, while between morph interactions are settled by chromatic badges.

### Morph and bib frequency variation

We found a frequency bias towards yellow males, and an absence of orange males in the populations we sampled; orange has also been consistently rare at Yathong (McDiarmid et al., 2017; Olsson, Healey, et al., 2007; Rollings et al., 2017). The blue male morph has increased at Yathong (McDiarmid et al., 2017; Olsson, Healey, et al., 2007; Rollings et al., 2017), and is now as frequent as the red morph. One of the conspicuous differences between our population and those previously described is the non-random expression of the bib across morphs. We find that bibs are extremely rare in red males, but randomly distributed in blue and yellow males, this contrasts with the Yathong population where bibs are randomly distributed across yellow and red males (Healey & Olsson, 2009). Non-random trait associations often arise from linked loci, but break down unless maintained by strong correlational selection (Sinervo & Svensson, 2002). Our data suggest divergent genetic architecture and selective pressures on head and bib traits of males in different populations. At Yathong, red males frequently have bibs, and similar to our finding, bibbed males were in relatively good condition (Healey & Olsson, 2009). This population difference may reflect stronger competitive selection on red males at Ngarkat, consistent with the tendency for blue males to win contests against red males, potentially rendering the additional costs of bib expression unsustainable for red males in this population.

### Distribution

Evidently there are wide-scale variations in morph frequencies and presence; however we also detected local patterns in the distribution of males. Blue males were more likely to have a neighbour that was red, and much less likely to have a neighbour that was yellow, but the distance between reds and blues and their neighbours was larger than yellows and their neighbours. We were unable however, to detect a dominance hierarchy across the male morphs in contest experiments. The closer proximity of yellows to each other, supports the findings of previous work that suggested that yellow males are less likely to engage in territorial competition, having fewer perches in their territories consistent with a cryptic sneaker strategy (Healey et al., 2007; Olsson, Healey, et al., 2007). Of the three intra-morph contest types, yellow males had the lowest aggression scores. Yellow males also had low aggression when interacting with blues and to a lesser extent reds. On one hand, we might expect monomorphic clusters of males to occur if some male morphs engage in passive or altruistic behaviours, for example black headed Gouldian finches are much more physiologically tolerant of a high frequency of other black headed males than the aggressive red headed finches are (Pryke et al., 2007). However, monomorphic clusters should always be unstable and liable to invasion from rare alternatives. The instability of monomorphic aggregations has been hinted at in *C. pictus* where releases of triads of painted dragons either comprised of a monomorphic trio of males and a female, or a polymorphic trio and a female, resulted in more subsequent recaptures of polymorphic combinations than monomorphic ones (Healey et al., 2007). This result could have been driven by an increased impetus to disperse, as we have shown aggression within morphs is greater than between.

### Condition dependence

Head dimensions are important estimators of bite strength in lizards (Herrel et al., 2001), and we found head dimensions were positively allometric. This pattern conforms to trait expectations in which larger males derive a greater fitness benefit from greater trait expression, consistent with condition-dependence (Fromhage & Kokko, 2014). Interestingly, we did not detect a difference in either the allometry of head dimensions among morphs or the size of the male morphs. Instead, the pattern for the allometry of head dimensions suggests that all male head morphs follow the same allometric trajectory. We had little power to detect an influence of bib expression on dominance between males, however, within yellow and blue males, those with bibs had a steeper allometry of head depth than those without, and a tendency to be heavier for their size. This pattern is accompanied by evidence for condition dependence in the bibs - bibbed males were in better condition, and the area of the yellow bib itself was related to condition and was positively allometric; previously colour intensity has been related to condition (McDiarmid et al., 2017). These patters share some commonality with findings from other populations, reinforcing the notion that bibs are indicative of higher condition (Healey & Olsson, 2009; McDiarmid et al., 2017; Olsson, Healey, et al., 2009). At Yathong there were bib by head morph interactions (Healey & Olsson, 2009), but those were between yellow and red males, whereas here we document only variation between blue and yellow males. This serves to highlight the significance of the non-random association between head-morph and bib expression.

### ‘Blue’ males

Blue males are described in the literature as ‘body colour’ males, since most males have a blue body colour and the distinguishing feature of blue males is that they lack red, yellow or orange (Baker, unpublished; Friesen et al., 2017; Friesen et al., 2021; Rollings et al., 2017). In our data we classified males as ‘no colour’ or ‘blue’ prior to any descriptions in the literature (i.e. 2003). We find a significant deficit of small blue males in the population however this deficit could be accounted for if the small males which we recorded with ‘none’ for head colour were putative blue males, that were not expressing the blue pigment. The suggestion from the data is that within the blue males there is potentially size- or age-dependent conditional expression of the blue head colour. There is no similar size-dependent conditionality in the expression of male red and yellow pigments. This would represent a novel conditional strategy within the alternative strategies of this lizard. Interestingly, in all male morphs of *C. pictus* the expression of blue colouration is body-size dependent, whereas red and green pigmentation is not (Baker, unpublished). Body size seems the most likely cue for the conditional (Hazel et al., 2004; Hazel et al., 1990) (as opposed to condition-dependent) expression of a trait in these lizards since we found that fights were on-the-whole won by the larger male even in size matched contests. The size dependence of blue expression may involve a developmental threshold hormone titre, or iridophore organisation, rather than the resource-dependent pigment deposition underlying red and yellow coloration (Olsson, Healey, & Astheimer, 2007). This distinction may explain the unique physiological profile of blue males compared to other morphs (Friesen et al., 2017; Rollings et al., 2017). If blue males evolved subsequent to red and yellow in *C. pictus*, the size-dependent expression of the head colour could have facilitated the invasion of this morph. Size is an important factor in fight outcome – hence a novel morph that was cryptic when small might invade more successfully than a novel morph that advertised its presence before it could win aggressive interactions. Consistent with this, putative blue males with no head colour were found close to other males. A genomic interrogation of the spatial and temporal pattern of morph evolution in *C. pictus* would shed light on this question (Küpper et al., 2016; Rankin et al., 2016).

### Aggression within and between morphs

Our finding that intra-morph aggression is heightened compared to inter-morph aggression, and that within, but not between morph interactions are resolved through differences in body size, provides convincing evidence that head colouration acts as a ‘badge of status’ that determines a behavioural strategy and allows rapid contest resolution with limited escalation (Maynard-Smith, 1982; Parker, 1974). This supports manipulative experiments in this species that showed that when males were unable to determine the morph of their opponent, that fights escalated (Healey & Olsson, 2009; Healey et al., 2007). Similar to Gouldian finches (Pryke & Griffith, 2006) we also found that interactions between the red-headed morphs tended to escalate; the higher testosterone titres in red males (Olsson, Healey, & Astheimer, 2007) are likely to account for these high intra-morph levels of aggression. Our data show contrast between attacking behaviour and displays in within vs between morph aggressive interactions. While lateral displays were common to both contest types, head-bobs increased in interactions between morphs in the opposite pattern to attacks. This supports the notion that chromatic signals resolve aggression between morphs, but not within. Elevated levels of aggression within compared to between morphs supports the idea that novel colour morphs might arise through negative frequency dependent benefits to novel colour variation (Seehausen & Schluter, 2004). For example, where dominant males are selected to defend their resources from strategists that they encounter most frequently, novel phenotypes could attract less attention from dominant males, providing a toe-hold for the evolution of a new morph. We did not stage contests with blue males that were in the small size category, and so whether small blue males bypass aggression from other males remains to be seen. The way lizards assess rivals, both visually and physically might explain the profusion of male morphs in some lizards including *C. pictus*.

## Conclusion

We document the morphology and behaviour of an understudied male morph in a system that is otherwise very well characterised. Our finding of non-random trait associations between head morph and bib, that is strikingly different to Yathong, suggest that there is divergence in the genetic architecture of morph expression between these populations. This difference suggests that there is variation in the correlational selection acting on head colours and bib expression and male condition. The non-random association also suggests that the superior condition of bibbed males is attributable to their genotype, rather than a product of it, as is the case in a conventional condition-dependent trait. Our data clearly show that badges reduce between morph aggression, and that within morph aggression is settled in a condition-dependent way. Hence selection is acting strongly within morphs, presumably dependent on their frequency, such that divergence between morphs in competitive ability could hitchhike on drivers of within morph competition. Finally, blue males, which are already recognised as physiologically unusual, appear to have novel conditional, size-dependent expression of head colouration, that may allow them to invade yellow-red dominated populations. This differs from the recent suggestion of territory-dependent conditional morph expression in side-blotched lizards (Corl et al., 2026), but, if supported, would constitute a novel conditional strategy within the suite of genetic strategies seen in this species.

## Acknowledgements

We thank BG & PR LeBas for field assistance and accommodation. NRL was funded by a NERC Fellowship.

